# Meta all the way down: An overview of neuroimaging meta-analyses

**DOI:** 10.1101/2025.01.21.634094

**Authors:** Kendra Oudyk, Jérôme Dockès, Julio Peraza, James Kent, Mohammad Torabi, Michelle Wang, Brent McPherson, Niusha Mirhakimi, Alejandro de la Vega, Angela R. Laird, JB Poline

## Abstract

Meta-analyses are invaluable tools for navigating the rapidly expanding scientific literature. Given their high value, ensuring the quality of meta-analyses is paramount. We conducted a multifaceted overview, examining each step in a manual neuroimaging meta-analysis on a large scale. We used four novel datasets comprising over 14,000 papers, including fMRI meta-analyses, fMRI studies, studies included in meta-analyses, and studies associated with image data on NeuroVault.

Regarding successes, two-thirds of meta-analyses stated that they followed PRISMA guidelines, and 65% included a flowchart describing their inclusion process.

We point out several areas for improvement. Pre-registration was fairly rare (20%), and only half listed their exact search strategy. There could be a location bias in which papers are included, and many did not include enough studies to be robust against publication bias (68% of meta analyses have less than 30 studies included).

We also offer ideas for future directions. As image based meta-analysis is the gold standard, we have indicated which topics have the most image data available. The potential redundancy of topics can be visualized in our paper, and we recommend future meta-analyses be in conversation with past ones by citing and discussing previous similar work. By addressing these findings, the neuroimaging community can collectively improve the field of neuroimaging meta-analyses.

## Introduction

The need to synthesize the scientific literature has never been more pressing, and meta-analyses and reviews are essential tools for this task. Meta-analyses offer systematic and robust methods for quantitatively summarizing previous research and for identifying novel findings (Tshitoyan et al., 2019). They are the strongest level of evidence in medical research (*Oxford Centre for Evidence-Based Medicine*, n.d.). Further, they are particularly important in fields like functional magnetic resonance imaging (fMRI), where individual studies tend to have low power due to the high cost of gathering data from many participants (Button et al., 2013). Meta-analysis offers a way to combine the results of many studies to increase power.

While neuroimaging meta-analysis is a growing field, with active work in both methods and applications, there has not yet been an overview of fMRI meta-analyses (though see Yeung et al., 2023 for an overview of meta-analyses that specifically use the BrainMap software). Due to their high regard in terms of evidence, meta-analyses deserve high scrutiny.

In the present study, our goal was to provide an overview of the field, pointing out potential issues, successes, and ways forward to help researchers generate more rigorous knowledge about brain structure and function. In this case, the term ‘overview’ has a specific meaning: it is a quantitative summary of reviews or meta-analyses (Gates et al., 2022; Higgins & Cochrane Collaboration, 2019). We did not perform a systematic review because this work was exploratory. Here, we have conducted an overview of published neuroimaging meta-analyses, which we present in terms of five main steps:

1. **Planning.** First, we aimed to help researchers identify topics with sufficient data for adequately powered image-based meta-analyses by showing which topics have the most image data. Next, we explored a novel method for investigating redundancy in meta-analyses, that is, when more than one meta-analysis is done on the same topic. Another important part of planning a meta-analysis is designing the study around rigorous guidelines. We showed which guidelines are being used most often in neuroimaging meta-analyses.
2. **Pre-registration.** It is recommended to pre-register the hypotheses and protocol by writing them out and sharing them publicly. This is important for meta-analyses, whose results are sensitive to researcher choices such as inclusion criteria. We asked how many meta-analyses have been pre-registered, and where. Pre-registration is also supposed to make it easier to publish null results, and we investigated whether pre-registration has been successful in this regard.
3. **Data collection.** A key step in neuroimaging meta-analysis publications is reporting publication databases and keywords used in literature searches (Rethlefsen et al., 2021). We summarized how meta-analyses reported their literature searches. Specifically, we examined the replicability of the search strategy, as well as the number of databases searched and whether that predicted the number of papers included. These factors are key for the inclusion process to be transparent, but it should also be free of bias. We explored the possibility of gender and/or location bias in the corpus of included papers. It is also important to include enough papers in the meta-analyses. We examined the adequacy of statistical power for primary studies included in meta-analyses, as well as for the meta-analyses themselves.
4. **Analysis.** Various methods have been proposed for performing neuroimaging meta-analyses. They generally fall into two main categories: image-based and coordinate-based meta-analyses. Image-based meta-analyses are preferred because they summarize results across brain images and retain un-thresholded information; in contrast, coordinate-based methods use the coordinates reported in published papers, which contain less information than the full results images. We reported how often different methods were used, considering which methods are preferred.
5. **Reporting and publishing.** The process of including/excluding papers is key for the results of a meta-analysis, and so it is important that these steps are reported transparently. We assessed the prevalence of figures like flow diagrams that show the inclusion process. Next, we looked at the step of publishing results, which can determine the impact of a meta-analysis. Specifically, we examine where meta-analyses are published, those journals’ impact factors, and whether journal impact factor relates to the number of citations in neuroimaging meta-analyses.

To answer the above questions, we searched the literature and identified four sets of scientific articles (see the Methods section for details): (1) 761 fMRI meta-analyses, (2) 36,766 fMRI papers, (3) 2,142 papers included in our set of open-access fMRI meta-analyses, and (4) 1,395 papers associated with results images stored on NeuroVault (Gorgolewski et al., 2015). Note that the set of meta-analyses included coordinate-based and image-based meta-analyses, as well as large-scale automated meta-analyses (e.g., NeuroSynth).

Our datasets were collected from PubMed and PubMed Central using <u>pubget</u> (Dockès et al., 2023), a tool for automatically gathering academic papers and metadata. Relevant information was extracted manually and automatically using labelbuddy and pubextract, respectively. These tools are part of the *litmining* ecosystem for collecting data and performing text mining on the scientific literature (Dockès et al., 2023), which make meta-research more accessible, efficient, and reproducible.

In the following section, we describe each meta-analysis step and subquestion, in terms of their specific background, methods, and results. We follow this section with a general discussion, and conclude by describing the data collection process.

The details and parameters of the methods used can be found in the code, shared at github.com/neurodatascience/review-neuro-meta-analyses. All of the data for this study is available at <u>osf.io/d2qbh</u>/ and the annotations can be found at github.com/litmining/labelbuddy-annotations/.

## Brief methods and results for each step

### Step 1. Planning

#### Which topics have sufficient data for a high-powered image-based meta-analysis?

A meta-analysis should consolidate, address, or find a gap in the literature, and it is important that a topic has enough data to support a well-powered meta-analysis. Further, one should aim to use the gold standard method, where possible; in neuroimaging, this means trying to focus on a topic with enough brain images available to do an image-based meta-analysis (IBMA). IBMAs use unthresholded brain images as opposed to coordinate-based meta-analyses (CBMAs), which use brain coordinates reported in journal articles. There is more information in a full brain image, so IBMAs are considered the best method (Salimi-Khorshidi et al., 2009). Note that CBMAs can be useful, but we are suggesting that authors use IBMA if possible.

To gather images for an IBMA, authors may contact the authors of those studies and request them, or they can obtain them from NeuroVault, an online public repository of unthresholded statistical maps (Gorgolewski et al., 2015). In this section, we aimed to identify topics that have the most data on NeuroVault, to help guide future meta-analyses.

In order to determine the topics in the fMRI field, we used non-negative matrix factorization (in Scikit Learn; Pedregosa et al., 2011) to decompose the matrix of term-frequency - inverse-document-frequency (TFIDF) values from the abstracts of 36,766 fMRI papers, with 10,928 terms, into 80 topics. We labeled each topic with the term that explained the most variance.

The process of getting topics in this way can be fully automated; however, some of the topics did not make sense from the perspective of cognitive neuroscience. For example, one topic had “set” as the highest-loading term; this does not sound like a topic compared to “autism” or “memory”. Therefore, we opted for a semi-automated approach where we extracted topics, then noted any terms that were not topic-related, added those terms to a list of ‘stop-words’, and removed those stop words before running the topic modeling again (stop words are available in the code for this project at https://github.com/neurodatascience/review-neuro-meta-analyses/blob/main/scripts/0_data-collection/4_fmri-papers/data/1_input_stop-words.txt.

To determine how many papers belonged to each topic (with overlap), we thresholded the normalized loading of individual papers onto the topic at 0.9 out of 1.0 (an arbitrary choice). Then we counted the number of papers in each topic in a) our set of participant-level fMRI papers, and b) our set of NeuroVault papers.

Figure 1 shows the correlation between the approximate number of fMRI papers versus NeuroVault papers (Spearman’s R = 0.68, p < 0.01, 95% CI = 0.54-0.78). We labeled each topic with the word with the highest weight in that topic. There is a strong correlation between these variables, with larger fMRI topics having more data on NeuroVault. Topics towards the right of the figure have more data on NeuroVault, and could be good candidates for future meta-analyses. The topics with the most image data are decision, reward, cognition, language, and memory.

**Figure 1.**
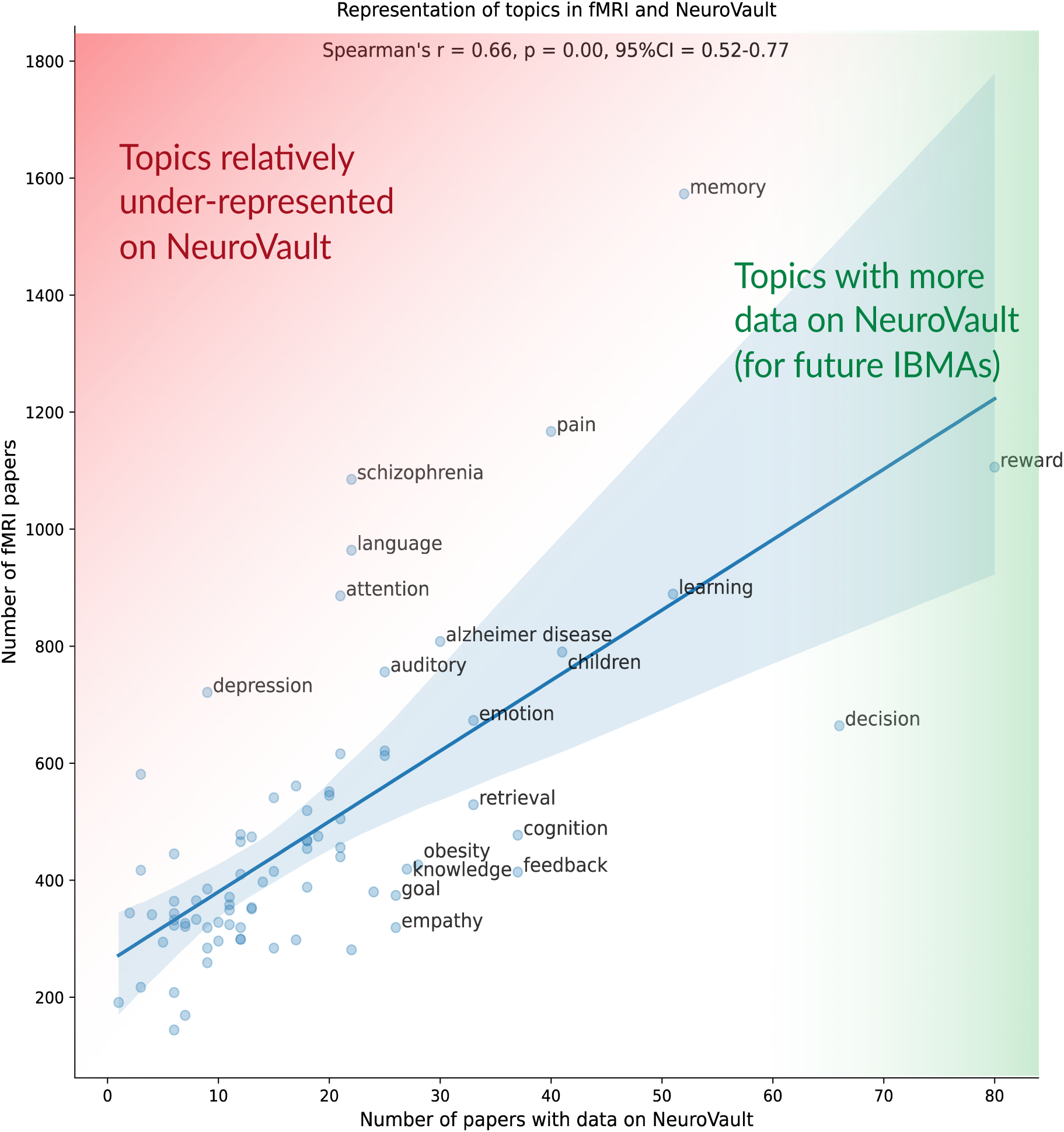
The number of papers with data on NeuroVault, compared to the number of fMRI papers for each topic. Topics towards the right of the graph have more data on NeuroVault. Topics towards the top left are relatively underrepresented on NeuroVault.

While image-based meta-analysis methods are preferred, coordinate-based methods used far more often (see Figure 6H). We suggest that authors of future image-based meta-analyses use Figure 1 to identify topics for which there may be sufficient power. This figure also shows which topics are relatively underrepresented on NeuroVault, and we suggest that authors with such data upload their results images onto NeuroVault.

#### Have authors considered previous meta-analyses on similar topics?

After selecting a topic, one must consider previous research, particularly previous meta-analyses on that topic. It has been suggested that redundancy is an issue in meta-analyses. A redundant meta-analysis is one where there have been previous meta-analyses on the same topic with a similar set of papers. Siontis and colleagues (2013) show that most meta-analyses (66%) have redundant meta-analyses published later; here, redundancy refers to having multiple meta-analyses on the same topic. Ioannidis (2016) shows that about 27% of meta-analyses overlap substantially with previous meta-analyses. He concludes that this is wasteful and unnecessary.

On the other hand, updated meta-analyses are needed at regular intervals. Shojania and colleagues (2007) estimated that quantitative systematic reviews can be considered out of date after a median of 5.5 years (and some were out-of-date at the time of publication). These authors and others advocate for *updating*, rather than re-doing, meta-analyses and systematic reviews on the same topic (Higgins & Cochrane Collaboration, 2019).

Conversely, one could argue that it is useful to replicate past research. The problem then occurs when previous research is replicated without being in conversation with the replicated research. Only 20% of meta-analyses discuss previous meta-analyses on the same topic (Helfer et al., 2015), so it is unclear whether teams are aware of previous work and to what extent their work might overlap.

Thus, here we describe the extent of redundancy in meta-analyses, without judging replications as good or bad. All of this previous research was done using manual methods, but we aimed to automate this process as much as possible. We addressed the question of redundancy with two approaches: first, we compared the number of papers in fMRI with the number of meta-analyses for different topics, and second, we described how many overlapping papers there are between meta-analyses on the same topic.

We used the same topic model as was used in the previous section, applying the weights to the set of meta-analyses. Figure 2A shows the relationship between the number of fMRI papers and the number of meta-analyses for each topic. There is a moderate correlation (Spearman’s R = 0.60, p < 0.01, 95% CI = 0.54-0.78). At the upper-left corner of the graph are topics that have many fMRI papers but few meta-analyses; this points to what could be meta-analyzed next. On the other hand, at the lower-right corner are topics that have few fMRI papers and many meta-analyses. These are topics that may have been well-covered by meta-analyses; one should consider how recent is the last meta-analysis, and whether another one is needed.

**Figure 2.**
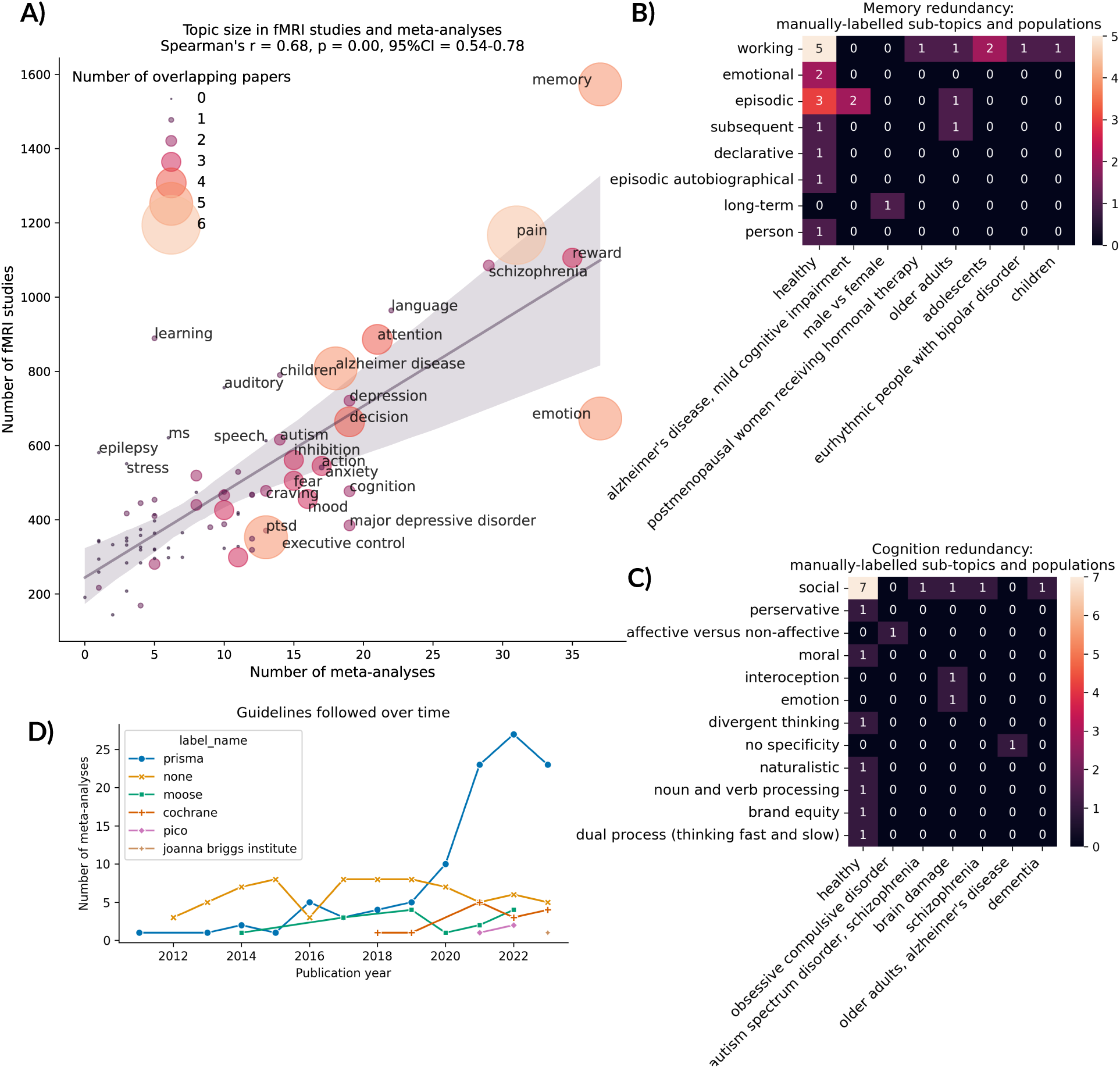
Considerations when planning a meta-analysis (redundancy and guidelines). A) Number of meta-analyses versus the number of fMRI papers on each topic, with the point size representing the number of overlapping papers in the topic (number of papers in more than one meta-analysis). B) and C) A closer look at the subtopics and populations examined in meta-analyses on the topics of Memory and Cognition. D) Guidelines followed in neuroimaging meta-analyses over time.

The topics in this graph are very broad. For example, there might be two very different meta-analyses on “memory”; one could focus on working memory in Parkinson’s disease, and the other could focus on episodic memory in healthy controls. These would not be redundant if one considers the subtopics and populations being studied. To determine how much Figure 2A can tell us about redundancy at a more fine-grained level, we manually inspected two topics with the same number of meta-analyses (22) but widely different numbers of fMRI papers: memory and cognition. We manually labeled the populations (e.g., Alzheimer’s, healthy control) and the sub-topic (e.g., working memory, social cognition). We measured redundancy as the number of meta-analyses with the same subtopic and population. We predicted that cognition meta-analyses would show more redundancy compared to memory, since there are fewer participant-level fMRI papers on “cognition” for the same number of meta-analyses.

Figures 2B and 2C show the results of this evaluation. For the memory meta-analyses, there were 12 papers with another paper on the same sub-topic and population (see Figure 2B). For the cognition meta-analyses, there were 7 (see Figure 2C). While we did not perform a statistical test on so little data, these numbers did not follow our prediction that cognition would be more redundant than memory. Therefore, this automated method of evaluating redundancy (i.e., Figure 2A) needs to be further refined.

#### Which meta-analysis guidelines are being used?

Meta-analyses are considered the highest form of evidence, thus, rigorous standards have been developed. Here, we briefly review these standards and quantify their use in neuroimaging meta-analyses.

In 1996, guidelines for reviews and meta-analyses became more standardized. A group of epidemiologists met and laid out a set of guidelines at the Quality of Reporting of Meta-analyses (QUOROM) conference (Moher et al., 1999). In 2009, these were replaced by the Preferred Reporting Items for Systematic reviews and Meta-analyses (PRISMA; Liberati et al., 2009), and which were updated again in 2020 (Page et al., 2021).

PRISMA applies primarily to biomedical fields, especially randomized control trials. However, the guidelines have also been adapted and used in other fields, such as observational studies (Meta-analysis Of Observational Studies in Epidemiology, MOOSE; Brooke et al., 2021) and nursing (Joanna Briggs Institute, JBI; Santos et al., 2018). The Cochrane Collaboration emerged in 1993; it is a community that conducts and archives high-quality systematic reviews and meta-analyses in the medical sciences. Cochrane papers are supposed to meet the PRISMA guidelines.

While the PRISMA guidelines focus on reporting of study selection processes (i.e., applying inclusion/exclusion criteria), Cochrane has an additional standard for ensuring more-broad methodological rigor: the Methodological Expectations of Cochrane Intervention Reviews (MECIR; (*MECIR Manual | Cochrane Community*, 2023). Further, the Cochrane Handbook suggests defining the scope and inclusion/exclusion guidelines based on the Population, Interventions, Comparators, and Outcomes (PICO; Higgins & Cochrane Collaboration, 2019). Essentially, Cochrane is a community/archive, whereas PRISMA, MOOSE, JBI, PICO, and MECIR are checklists/guidelines.

In this section, we asked which guidelines are being applied in neuroimaging meta-analyses, and how often. We manually annotated which guidelines authors said they followed in the open (full-text) fMRI meta-analyses in our dataset. We found that the largest group of papers used PRISMA guidelines (54.6%), followed by papers using no guidelines (37.2%), and then MOOSE (8.2%), Cochrane (7.1%), PICO (2.5%), and JBI (0.5%). Further, we found that the number of papers using PRISMA guidelines is increasing over time, which represents a success in our field (see Figure 2D).

This increase may reflect the heightened emphasis on rigor and transparency in research. Following standards promises to decrease bias and increase reliability in research.

### Step 2. Pre-registration

#### How many meta-analyses are pre-registered, and where?

Pre-registration of scientific studies is meant to address the problem of publication bias, where studies with null results are less likely to be submitted and accepted for publication (Easterbrook et al., 1991; Rosenthal, 1979; Thornton & Lee, 2000), as well as the problem of researcher degrees of freedom, where researchers increase the likelihood of false-positive findings when they try multiple analysis methods until they find the results that they anticipated (Hardwicke & Wagenmakers, 2023).

These problems are present in the broader field of primary studies, as well as in reviews and meta-analyses. Both the Cochrane Handbook (Higgins & Cochrane Collaboration, 2019) and the PRISMA guidelines (Page et al., 2021) recommend pre-registration of protocols for meta-analyses and reviews.

First, we asked how many neuroimaging meta-analyses were pre-registered, and where the pre-registrations were archived. We annotated this information manually in 201 open (full-text) meta-analyses. We found that 20.4% of meta-analyses were pre-registered. As can be seen in Figure 3A, 15.4% were pre-registered in the International prospective register of systematic reviews (PROSPERO; Moher et al., 2014), 3.5% in the Open Science Framework (OSF; Foster & Deardorff, 2017), and 1.5% on the International Platform of Registered Systematic Review and Meta-analysis Protocols (INPLASY; Canellas et al., 2022). Note that the latter is not free, while the first two are.

**Figure 3.**
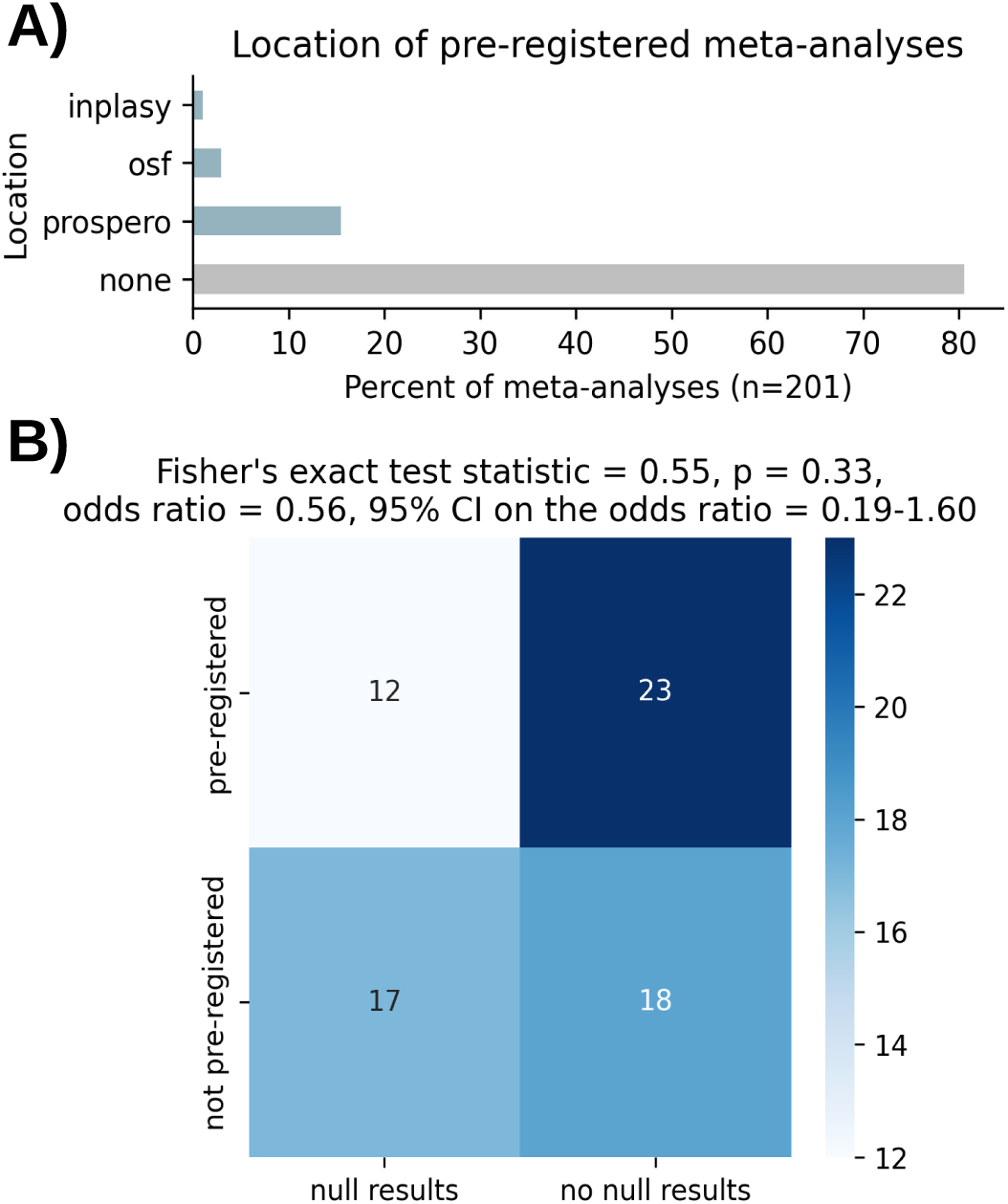
A) Location of pre-registered neuroimaging meta-analyses. B) Contingency table for whether meta-analyses are pre-registered and whether they report null results.

#### Are pre-registered meta-analyses more likely to have null results?

Next, we wanted to explore whether pre-registered meta-analyses have more null results. One of the goals of pre-registration is to address the problem where it is more difficult to publish null results. This is because of publication bias, and it results in the file drawer effect, where null results are ‘put in the file drawer’ instead of being published. Previous research has shown that 50-60% of pre-registered studies have null results, compared to 5-20% of non-pre-registered studies (Allen & Mehler, 2019). Thus, we predicted that the pre-registered studies would have more null results.

To address this question, we manually labeled whether or not studies reported null results in all of the pre-registered meta-analyses (n = 35) and an equal number of non-pre-registered meta-analyses (n = 35). This information allowed us to create a contingency table, counting papers in groups defined by whether they were pre-registered and whether they had null results. Using Fisher’s exact test, we found no significant differences between these groups (probability = 0.55, p = 0.33, odds ratio = 0.56, 95% CI on odds ratio = 0.19-1.60); we did not find that the pre-registered studies had more null results.

This analysis had a limited sample size due to the fact that only 35 of the open meta-analyses were pre-registered, and therefore may lack statistical power and give biased results. We therefore do not conclude in either direction (whether pre-registration is helpful for combatting the file drawer problem, or not).

### Step 3. Data collection

#### How replicable and comprehensive are the searches in meta-analyses?

Well-conducted and rigorous meta-analyses should strive to be comprehensive, including all relevant papers or a representative sample. Further, the search should be replicable so that the meta-analysis may be replicated and/or updated in the future. In recent years, there has been increasing guidance on how to search for papers and how to report the results of that search (Higgins & Cochrane Collaboration, 2019; Page et al., 2021).

First, we asked how reproducible the search results are in neuroimaging meta-analyses. We manually annotated the searches in 77 meta-analyses as a) having the exact search terms (i.e., text that could be copied into the database website to replicate and extend the search), or b) having a description of the search terms. We found that just over half (52%) of meta-analyses had exact search terms. While the number with exact searches appears to be increasing over time (Figure 4A), this presents significant room for improvement.

**Figure 4.**
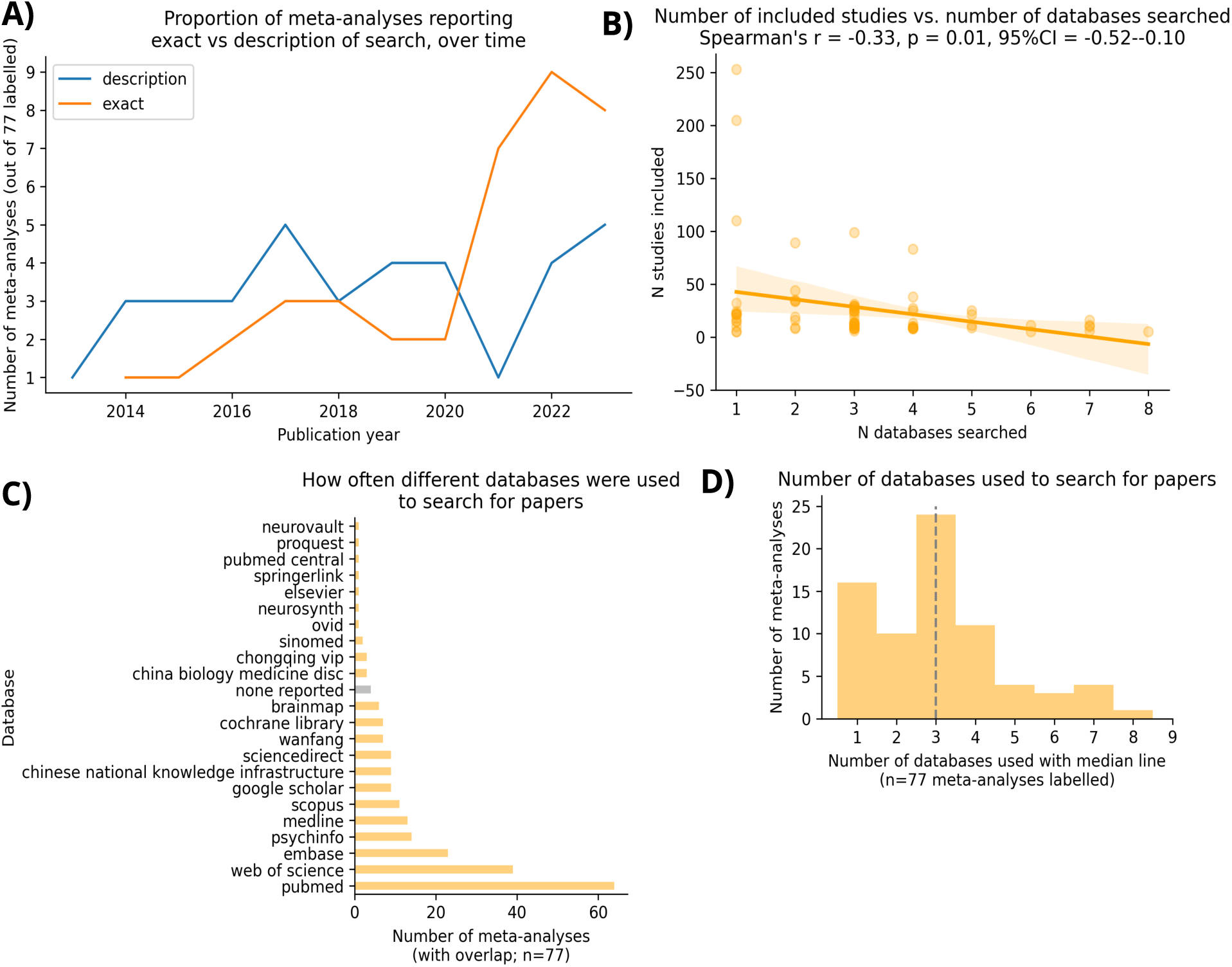
Aspects of data collection. A) Proportion of meta-analyses reporting exact versus description of search, over time. B) Relationship between the number of studies included versus the number of databases searched. C) How often different databases were used to search for papers. D) Number of databases searched for papers to include in meta-analyses.

Next, we asked how comprehensive are the searches in neuroimaging meta-analyses by looking at the number of databases searched. We also report how often different databases are used, as a guide for future meta-analyses. We manually annotated the databases searched in 77 meta-analyses. We found that the highest proportion of meta-analyses searched three different databases (range 1-8; Figure 4B). The most popular databases were PubMed, followed by Web of Science and EMBASE (Excerpta Medica Database; Figure 4C). We further wanted to know if searching more databases led to more studies being included in the meta-analyses. We found a slight negative correlation (Spearman’s R = −0.33, p = 0.01, 95% CI = −0.52 to −0.10), where searching more databases was related to including fewer studies (Figure 4D). It could be that, when meta-analysts find few papers to include, they proceed to check more databases to expand their pool of papers.

#### Is there gender and/or location bias in which papers are included in meta-analyses?

All humans have bias, and this bias can influence research in measurable ways. Regarding gender bias, Dworkin and colleagues (2020) demonstrated that women tend to be under-cited in the neuroimaging literature, particularly when the citing authors are men. Further, regarding racial bias, Bertolero and colleagues (2020) showed that White authors tend to cite other White authors more. In line with this, Liu and colleagues (2023) found that Non-White scientists receive fewer citations. Citations are an important currency in science (Carpenter et al., 2014), and such biases could have an impact on the careers of minority scientists. However, it is not known whether similar biases exist when papers are being selected for meta-analyses (cf., Rogers & Seaborn, 2023). This would have implications for whose work is being included in the crystallization of knowledge within a field, seeing as meta-analyses can be considered the highest form of evidence (*Oxford Centre for Evidence-Based Medicine*, n.d.).

To investigate whether there is gender bias in the inclusion process in neuroimaging meta-analyses, we tested the null hypothesis that the gender proportions in included papers equals the gender proportions in the broader fMRI field. We estimated author genders from author names, recognizing that this imposes a false binary on gender that may not reflect the lived experiences of authors. We took these automatically-extracted names for the authors of fMRI meta-analyses (n = 36,766) and the papers included in open-access fMRI meta-analyses (n=2,142), and looked for a match in the World Gender Name Dictionary 2.0 (Lax-Martinez et al., 2023). This dictionary includes more than 26 million records of given names, from 195 different countries and territories. It records the gender typically associated with each name. Following the precedent of Dworkin and colleagues (2020), we focused on the genders of the first and last authors, since these are typically the key authors of neuroimaging papers. Further, following this example, we visualized the results in terms of categories of the gender of first and last authors: Woman first author - Woman last author, Woman-Man, Man-Woman, and Man-Man. For each author-gender category, we tested the null hypothesis using binomial proportions, with a one-directional test. For example, we wanted to determine whether there are proportionally fewer papers with a woman first and last author included in meta-analyses, compared to the proportion of papers with a woman first and last author in the broader fMRI field. We corrected for familywise error (because we did four tests) using Bonferroni correction. We found no significant differences in the proportions of papers in the author-gender categories between included papers and the broader fMRI field. This is not a confirmation of a lack of bias, since we are not doing a test to prove the null hypothesis. However, the apparent lack of bias may result from the rigorous inclusion procedure involved in meta-analyses, a procedure that perhaps protects against gender bias.

Next we wanted to answer the question of whether there is location (geographical) bias in the meta-analysis inclusion process. We would define location bias as an increased likelihood of including papers from countries with more money for research. Thus, we tested the null hypothesis that the number of papers included in meta-analyses from each country was not related to the Research and Development Expenditure (measured as a percent of Gross Domestic Product) of each country. In order to extract locations from the first authors’ affiliations, we listed the unigrams (single words) and bigrams (pairs of words next to each other) of the text in the affiliation, and took the intersection of this set and a set of 41,002 world cities (*World Cities Database | Simplemaps.Com*, n.d.), also searching for the country if there was more than one city with the same name. Next, in each dataset, we calculated the proportion of fMRI papers included in each country (the number of papers included in meta-analyses, divided by the number of fMRI papers on PMC). We then computed the correlation between these proportions and the countries’ Research and Development Expenditure (obtained from Our World in Data; *Research & Development Spending as a Share of GDP*, n.d.). There was a small, significant correlation (Spearman’s R = 0.33, p = 0.03, 95% CI = 0.03-0.57; Figure 5). This small correlation indicates that there is a relationship between the Research and Development Expenditure of countries and proportion of their studies that are included in meta-analyses. This may indicate location bias, e.g., where authors of meta-analyses are biased to include papers from countries with higher Research and Development Expenditure. However, there could be other interpretations of this finding. Perhaps countries with lower Research and Development Expenditure tend to collect data from fewer participants, and thus their papers are less likely to make it into a meta-analysis (meta-analysts sometimes exclude papers with too-few participants; Figure 6E).

**Figure 5.**
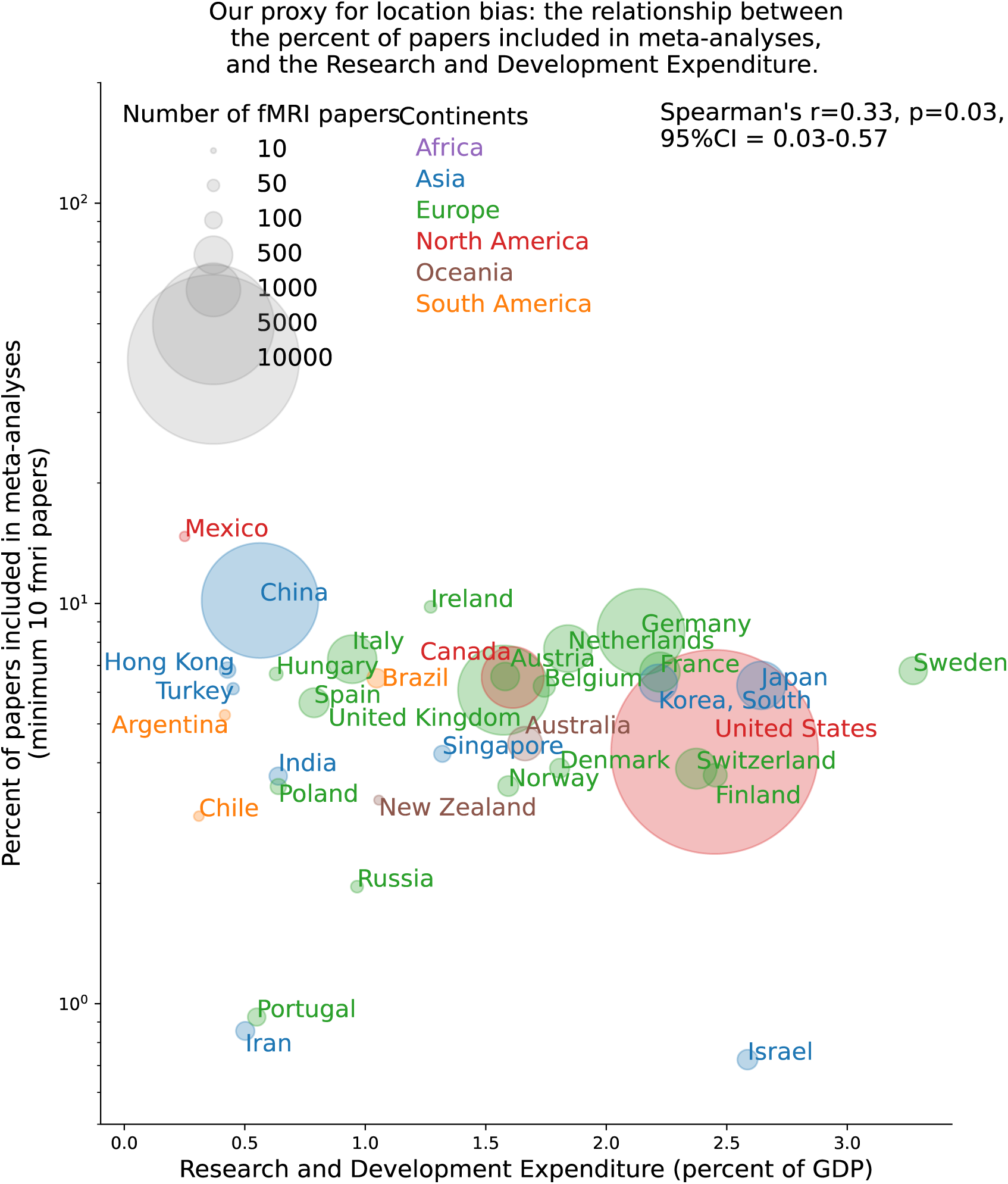
Our measure of location bias in data collection. Relationship between the Research and Development Expenditure of each country, compared to the percent of papers from that country that are included in meta-analyses. The size of the points represents the number of fMRI papers coming from that country, and the color of the country label represents the continent.

It is difficult to assess such biases in meta-research, since we have not directly observed the behaviors of meta-analysts. While we found no evidence of a gender bias in which papers are included in meta-analyses, we found some evidence of a location bias. It is important to mitigate a potential bias such as this, perhaps by having meta-analysts blind themselves to the author affiliations when deciding whether to include a paper in a meta-analysis.

#### Do meta-analyses include enough papers to be well-powered?

Concerns exist with reproducibility in science in general (e.g., Ioannidis, 2017), and it has been proposed that low power is one main cause of this problem (Button et al., 2013). In neuroimaging, Poldrack and colleagues (2017) demonstrated that we do not collect enough data to detect the effect sizes for the questions we ask (the usual effect sizes being around 0.5-1.5). Indeed, Marek and colleagues (2022) demonstrated that we need more individuals - an order of magnitude more - to detect stable effects. One proposed solution to this problem is meta-analysis (Wager et al., 2007). When results are pooled across multiple studies, statistical power is effectively increased. However, meta-analyses can have power problems as well. Eickhoff and colleagues (2016) provided the recommendation that a minimum of 17-20 experiments is needed to detect moderate effects, and later, Acar and colleagues (2018) suggested 30 experiments are needed to be robust in the face of publication bias. These studies specifically examined the method activation likelihood estimation (ALE; Eickhoff et al., 2009), a popular coordinate-based meta-analysis method (see Salo et al., 2023 for an explanation of such methods).

In order to investigate whether ALE meta-analyses have followed these recommendations, Yeung and colleagues (2023) reported the sample sizes in meta-analyses that used the BrainMap software (Fox & Lancaster, 2002). They found that about 20% included 17 or fewer studies. Here, we replicated and extended the results of Yeung and colleagues (2023) beyond BrainMap-based meta-analyses. To this end, we manually annotated the number of studies included in meta-analyses (including full texts of open-access meta-analyses and closed-access abstracts). Figure 6C shows a skewed distribution of the number of papers included in meta-analyses, with many meta-analyses including fewer papers. Further, while the median number of papers included has been steadily increasing over time (Figure 6B), it has not yet exceeded 30, the minimum proposed by Acar and colleagues (2018). In total, 68% of meta-analyses have fewer than 30 papers included.

**Figure 6.**
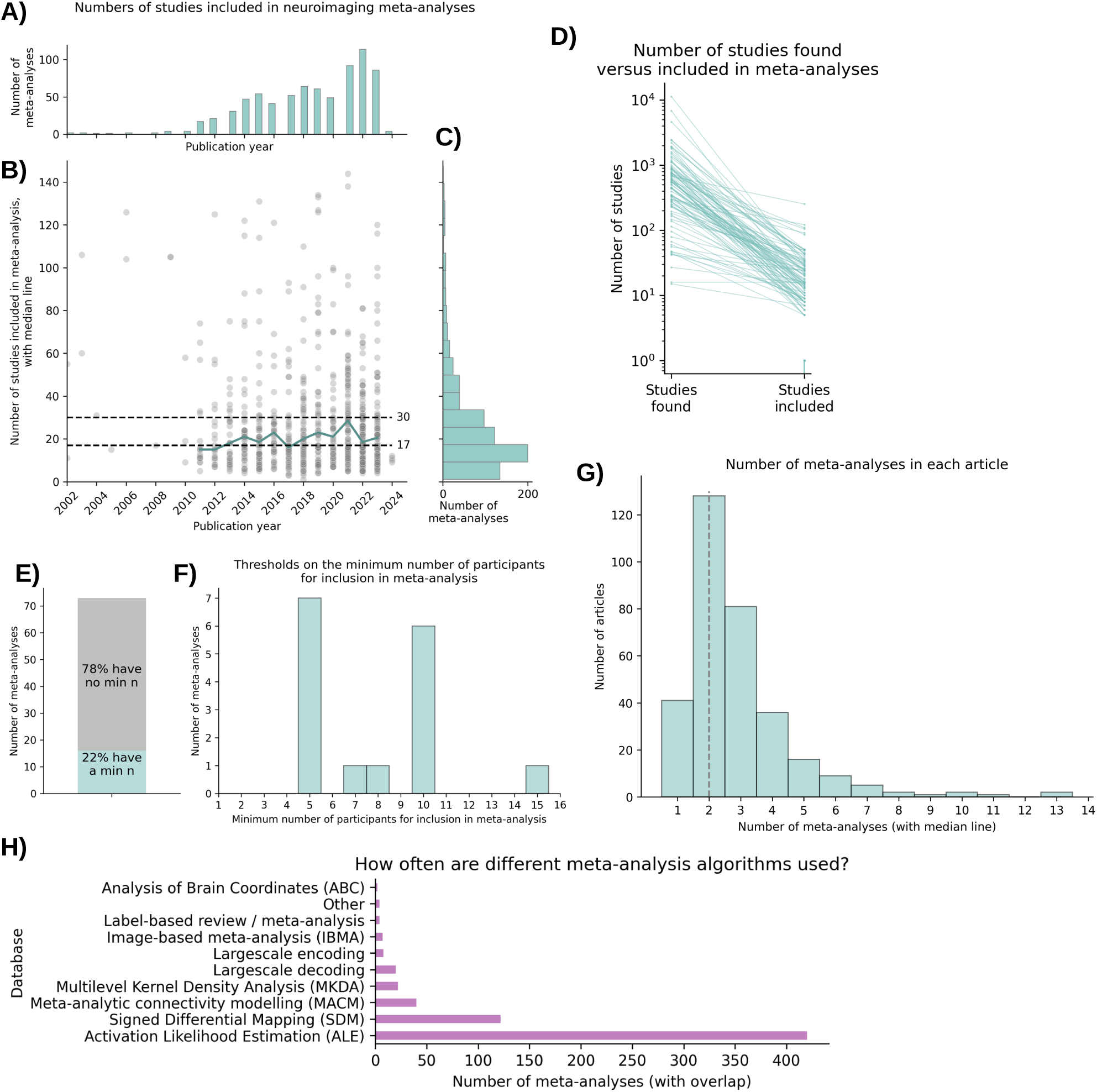
Aspects of power and meta-analysis algorithms. A) Number of meta-analyses published each year. B) Number of studies included in meta-analyses, over time. C) Distribution of the number of studies included in meta-analyses. D) Number of studies found versus number of studies included (one line per meta-analysis). E) How many meta-analyses use a threshold on the number of participants when deciding which papers to include. F) Thresholds on the number of participants for a paper to be included in a meta-analysis. G) Number of individual meta-analyses included in each paper. H) How often different meta-analysis algorithms are used.

While annotating the methods used in neuroimaging meta-analyses. We observed that many papers include more than one meta-analysis, and we decided to annotate it as well. The most common number of meta-analyses was two, with a range of one to 13 (see Figure 6G). This observation could raise concerns about family-wise error, since the meta-analyses within a paper are often performed on overlapping sets of primary studies. Future research using simulations could estimate the risk of increased false-positives when performing more than one meta-analysis on overlapping sets of data.

#### Do meta-analyses exclude papers with too few participants?

Another important consideration in meta-analyses is whether the included studies have enough power, that is, whether they include enough participants. As the saying goes, “Garbage in, garbage out”. This means that if the included studies are too noisy or are of poor quality, then the meta-analyses will also be noisy or poor quality (Sotola, 2022). Some suggest weighting the contribution of studies to the meta-analytic result according to their sample size, so that studies with low power contribute less to the meta-analytic result (Wager et al., 2007). This is normal practice in biomedical meta-analyses outside neuroimaing, but has yet to be made standard in this field.

We wanted to know whether meta-analysts are using a minimum threshold on the number of participants, and what those thresholds are. We manually annotated 77 studies and found that 22% have such a threshold (Figure 6E). The most common thresholds were five participants (seven papers had this threshold) and ten participants (six papers), with one paper for each of the thresholds seven, eight, and 15 participants (Figure 6F). It is unclear where these thresholds come from; none of these papers gave a citation or a reason for their choice in threshold. Today, an fMRI study with 5-10 participants would likely not be accepted into a peer-reviewed journal (one exception would be a dense sampling study with many data points per participant). Therefore, these thresholds might be considered low, and it may be concerning that three-quarters of meta-analyses do not have a threshold.

### Step 4. Analysis

Methods in neuroimaging meta-analyses is a rich and innovative field. There have been multiple methods and softwares created, some of which are argued to be better than others. For example, the class of image-based meta-analyses is preferred over the class of coordinate-based meta-analyses, because they preserve more information, whereas coordinates are lossy summaries. That being said, it can be more challenging to gather published image data. While coordinate data are readily available in the tables of fMRI publications, image data must either be gathered by contacting the authors of participant-level publications or by navigating NeuroVault, the online repository of unthresholded results images. How often are different methods used?

We wanted to know how often different meta-analysis methods are used, particularly regarding coordinate-versus image-based methods. To this end, we manually annotated all of the open full-text meta-analyses (n = 247), as well as the abstracts of closed-access meta-analyses (n = 652). As shown in Figure 6H, we found that the largest number of meta-analyses uses the activation likelihood estimation (ALE; 72.7%) method, followed by the signed differential mapping (SDM; 21.1%) method. On the other hand, very few papers used an image-based meta-analysis method (1.2 %). One of the goals of this paper was to make image-based methods more accessible, by making the data on NeuroVault more visible. We hope there will be more image-based meta-analyses in the future, perhaps on topics that we identified as having more data on NeuroVault (Figure 1).

### Step 5. Reporting and publishing

#### How transparently reported is the study selection process?

Over the past few decades, there has been a lot of effort to standardize and improve systematic reviews and meta-analyses. This is because they are considered the highest form of evidence, particularly in medical fields, where results can directly impact patients’ lives. Perhaps the most notable effort in this direction is the PRISMA guidelines (Moher et al., 2010). We wanted to determine to what extent neuroimaging meta-analyses align with the PRISMA standard. To do this in an automated way, we focused on whether meta-analyses included a flowchart figure, which outlines the process of including/excluding papers from the meta-analysis. It is important for this process to be transparent, objective, and reproducible in order to ensure that the meta-analysis results are not biased. We asked how often meta-analyses included a flowchart of the inclusion process. To answer this, we applied a heuristic method to automatically detect whether a meta-analysis contained a flowchart figure. The heuristic was that, for each paper, we looked through the figure descriptions for a set of keywords, including “flowchart”, “flow-chart”, and “PRISMA” (for a full list, see our open code). If a keyword was identified, the paper labeled as including a PRISMA-like flowchart. We validated this method by manually annotating 57 papers and then applied the heuristic method for automatic detection of a PRISMA-like flowchart on the same set of studies. Testing against held-out ground truth data, our heuristic method had an accuracy of 96.5%. We found that approximately two thirds of papers included a flowchart.

Figure 7A shows that the proportion of papers including a PRISMA-like flowchart has been increasing over time and is currently near 100%. This is an encouraging trend, as greater transparency can only improve the accuracy and reproducibility of meta-analyses.

**Figure 7.**
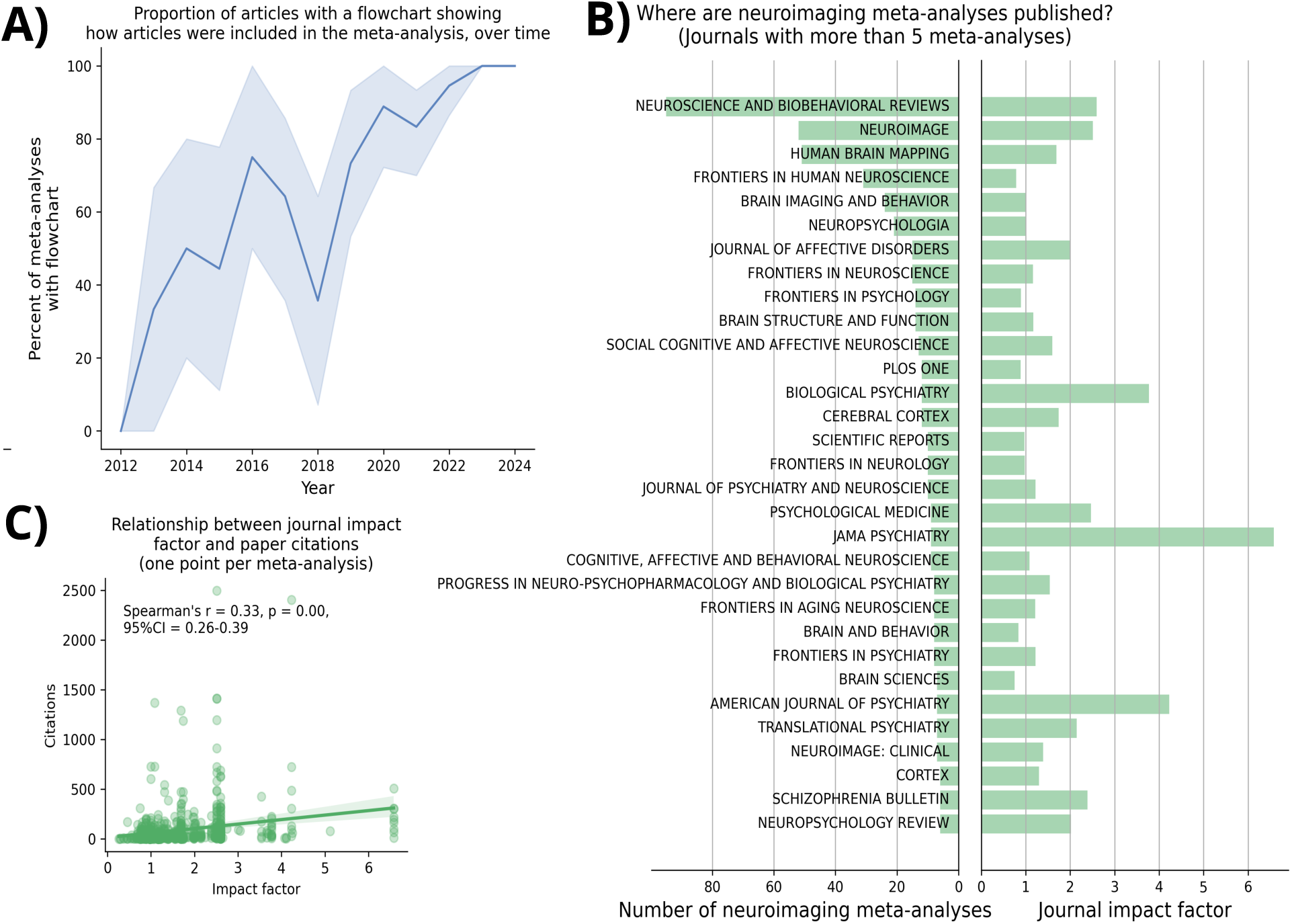
Aspects of reporting/publishing meta-analyses. A) Proportion of articles with a flow chart illustrating the inclusion process, over time. B) List of journals with more than 5 meta-analyses published in them. The right side shows the number of meta-analyses published in each journal, and the left side shows the impact factor. C) Relationship between the citation count of a meta-analysis and its journal’s impact factor.

#### Where should meta-analyses be published to have the greatest impact?

Academics need to publish to both communicate their work to their peers and to maintain and forward their careers. On the other hand, the focus on publication can lead to less-than-ideal practices in science (Smaldino & McElreath, 2016). Despite the pitfalls of focusing on publications, we feel it is an important enough step to be considered here. First, we wanted to know where meta-analyses are being published, and what is the impact factor of those journals. To this end, we used the journal meta-data extracted automatically by pubget, as well as impact factor data from the Scimajo Scientific Journal Ranking (*SJR : Scientific Journal Rankings*, n.d.). In Figure 7B, for the journals with at least 5 neuroimaging meta-analyses published in them, we plot the number of meta-analyses published (left) and the journal’s impact factor (right). Notably, several journals have the highest number of meta-analyses published in them (*Neuroscience and Biobehavioral Reviews*, *NeuroImage*, and *Human Brain Mapping*), whereas the rest of the journals have published few meta-analyses. Another observation from this figure is that the psychiatry-related journals have the largest impact factors. Authors concerned with impact factors and seeking to publish meta-analyses related to psychiatry could therefore target these journals.

#### Does journal impact actually predict citations?

A logical next step was to determine whether journal impact factor predicts the ‘impact’ of a meta-analysis paper, defined as the number of citations. We found a small correlation between journal impact factor and the number of citations of neuroimaging meta-analyses (Spearman’s R = 0.33, p < 0.01, 95% CI = 0.26-0.39; Figure 7C). Unfortunately we cannot interpret this relationship causally: one interpretation is that publishing in a higher-impact journal will lead to more citations; conversely, another interpretation is that better-quality meta-analyses both get accepted into higher-impact journals and also are cited more due to their quality. Note that another limitation is that we did not consider time as a variable in this comparison; older papers likely have more citations.

## Discussion

Meta-analyses can be considered the highest form of evidence. The purpose of this paper was to scrutinize the field of neuroimaging meta-analyses, focusing on each step in the process of conducting a meta-analysis. Here, we review the results in terms of successes, areas for improvement, and future directions.

### Successes

Regarding successes, there is increasing adherence to guidelines for meta-analyses, particularly the Preferred Reporting Items for Systematic reviews and Meta-Analyses (PRISMA; e.g., Page et al., 2021). About half of papers reported following these guidelines. Further, an increasing number of meta-analyses have a PRISMA-like inclusion flow chart, such that nearly all meta-analyses in the past few years have had one. Thus, the major successes that we observed are related to the increasing adherence to guidelines for meta-analyses. Continuing in this manner should improve the reliability and transparency of neuroimaging meta-analyses.

### Areas for improvement

While many studies used the PRISMA guidelines, a considerable proportion used no guidelines. Further, only 20% of meta-analyses were pre-registered; this is an important step for mitigating researcher degrees of freedom and biases. However, we did not find that pre-registration increased reporting of null results, as it was intended to do.

Once a meta-analysis has been pre-registered, the authors can begin searching for papers to include. This search should be thorough and replicable because it is highly recommended to update past meta-analyses rather than creating new meta-analyses on the same topic as previous ones (Higgins & Cochrane Collaboration, 2019). Just over half of searches were reported with the precise content and syntax used in the search, with the other half having only descriptions of the search strategy.

The next step is key: deciding which papers to include in the meta-analysis. Arguably, this is the step where researchers’ decisions have the greatest impact on the results, and it is important that it is unbiased. We found a positive relationship between a country’s proportion of papers included in meta-analyses and the percent of that country’s Gross Domestic Product (GDP) that is dedicated to Research and Development Expenditure. This could reflect a location bias, but there could be other explanations for this finding.

Another important aspect of which papers are included in the meta-analysis is power: both the power of the primary studies included, as well as the power of the meta-analysis itself. About a quarter of meta-analyses have such a threshold, with the most common threshold being five participants, followed by a second most common threshold of ten participants. This number appears to be arbitrary, and is too low if the goal is to only consider studies that would be of publication-quality today. Further, we found that many meta-analyses do not have sufficient power to be robust to publication bias.

Once there is a pool of studies to be included, the next step is to actually meta-analyze the studies. Some methods are recommended more highly; image-based meta-analyses are considered the gold standard method because they use the entire results images rather than only the reported coordinates of peak results. We found that image-based methods are seldom used and coordinate-based methods are more common, particularly activation likelihood estimation (ALE) and signed differential mapping (SDM).

### Future directions

Moving forward, we have several recommendations for future authors of meta-analyses. For those in the planning stage, we show that there is a lot of data available on NeuroVault for topics that are also larger in the broader fMRI field, particularly for the topics of “decision” and “reward”. We hope that Figure 1 will help future meta-analysts choose a topic with substantial image data on NeuroVault, enabling them to conduct an image-based meta-analysis, since IBMAs are the preferred method.

In order to help authors situate their topic appropriately, we explored a method for automatically evaluating meta-analytic redundancy, by comparing the number of primary studies to the number of meta-analyses for different topics in neuroimaging. However, closer manual inspection of some topics revealed that our automated method did not predict redundancy when considering more fine-grained subtopics and populations. Authors of future meta-analyses should consider past research on similar topics and populations when selecting a topic. Replicating a past meta-analysis is not necessarily bad, but authors should at least cite the meta-analyses being replicated. Seeing as many meta-analyses do not have sufficient power, authors could aim to extend those low-powered meta-analyses in order to solidify our knowledge in those fields.

Regarding location bias, future behavioral research could test whether country of origin is a (perhaps subconscious) factor in the process of deciding which papers to include in meta-analyses. Until we understand this phenomenon better, our recommendation is that meta-analysts should blind themselves to the country of origin when considering which studies to include in a meta-analysis.

Finally, there is the step of publication. We show that psychiatry-related journals have the greatest impact factor (*Biological Psychiatry*, *JAMA Psychiatry*, and the *American Journal of Psychiatry*). Further, we found a small positive relationship between journal impact factor and the number of citations for meta-analyses. Authors could consider these results when choosing where to submit their meta-analyses.

### Limitations

Although we present the most comprehensive overview of fMRI meta-analyses to date, there are several limitations. The principal limitation is that, while we retrieved the full texts of open-access publications indexed on PubMed Central, we only accessed the abstracts and meta-data of closed-access papers indexed on PubMed. This points back to a limitation of the pubget tool that we used to retrieve documents and meta-data. We were able to extend this tool so it could access abstracts from PubMed in addition to full-texts from PubMed Central, but ideally we would be able to retrieve full texts for all of the papers we examined. This is especially the case of the dataset of meta-analyses, in which we generated manual annotations that were not always possible when we only had the abstracts. For example, authors do not always list their methods in the abstract, so this could not be annotated when we only had the abstract. Another limitation of this study is the amount of work that was done manually. Many items were manually annotated in the meta-analyses, which limits the reproducibility and extendability of this study. However, we have shared the annotations at github.com/litmining/labelbuddy-annotations, so future research could aim to automate the extraction of features, such as which guidelines were used, which databases were searched, etc.

## Conclusion

This paper presents a comprehensive examination of fMRI meta-analyses. By assessing each step in a meta-analysis, we offer a holistic picture of the field and practical recommendations for future work. Authors should continue to follow PRISMA guidelines in terms of having an inclusion flow-chart. However, they can improve on the reporting of search terms in order to enable future extensions of their work. We highlight that few image-based meta-analyses are done, despite them being the gold standard method; authors may refer to this work to discover which topics may have sufficient data for a high-powered image-based meta-analysis. We address topics under debate, such as the usefulness of pre-registration and the level of power in meta-analyses. We hope that following these recommendations will improve the planning, performing, and publishing of neuroimaging meta-analyses, and that the field may continue to move towards more robust and reliable knowledge in neuroimaging.

## Methods

Here we describe our process of collecting the four datasets used in this paper. Details regarding automated data extraction may be found in the code at github.com/neurodatascience/review-neuro-meta-analyses and in Supplementary Figure 1. Details regarding manual data extraction may be found in the annotations repository at github.com/litmining/labelbuddy-annotations/tree/main/projects/neuro-meta-analyses and in Supplementary Figure 2.

### Collecting dataset 1) meta-analyses

We found a total of 940 papers, with 247 on PubMed central, and 693 on PubMed.

#### PubMed Central

We initially searched for open-access articles from PubMed Central using pubget (Dockès et al., 2023). Pubget is an open-access command-line utility to download papers from PubMed Central, and to extract metadata and full texts from the PMC API outputs. We used the following search: “(fMRI[Abstract] OR functional MRI[Abstract] OR functional magnetic resonance imaging[Abstract]) AND (meta-analy*[Title] OR meta analy*[Title])”.

#### PubMed

Pubget is limited to open-access papers, and a search on PubMed revealed that we were missing many papers that were not open access. Therefore, we searched for articles on the PubMed website and downloaded the list of resulting PubMed IDs. We then used those IDs as input to adjusted pubget code to automatically download the metadata and abstracts from the PubMed API. We used the following search on PubMed: “(fmri[Title/Abstract] OR functional MRI[Title/Abstract] OR functional magnetic resonance imaging [Title/Abstract]) AND (meta-analy*[Title/Abstract] OR meta analy*[Title/Abstract])”.

##### Inclusion criteria

Manual annotation was performed using labelbuddy (Dockès et al., 2023). We annotated the information of interest and which papers to exclude at the same time. To be included, papers had to:

- Be a meta-analysis of images/coordinates, not a traditional effect-size meta-analyses;
- Have humans as subjects;
- Be a published paper, not a preprint or a protocol;
- Be about the brain, not other organs;
- Be a meta-analysis, not about meta-analyses;
- Be a meta-analysis, not a review (though it could be both); and
- Be a meta-analysis, not use a previous meta-analysis as a seed for a participant-level study.

See Figure 8 for the number of papers excluded for each criteria.

**Figure 8.**
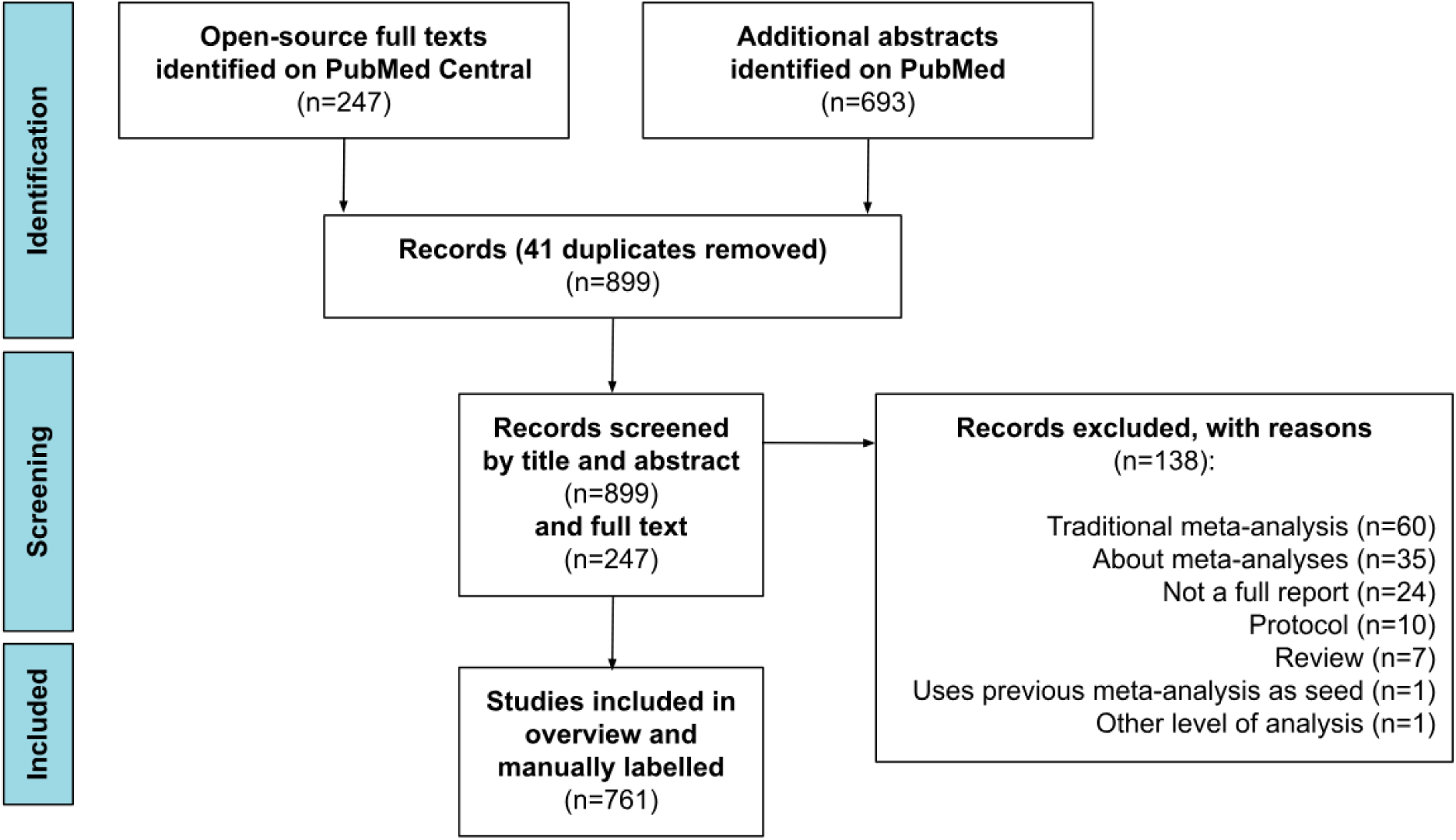
Flow chart outlining how data was collected and included/excluded.

### Collecting dataset 2) fMRI papers

We collected a dataset of abstracts and meta-data for fMRI papers from PubMed Central. We did this using pubget, without the option of restricting to open-access articles. We used the following query: “(fmri[Title/Abstract] OR functional MRI[Title/Abstract] OR functional magnetic resonance imaging [Title/Abstract]) AND (human)”.

### Collecting dataset 3) papers included in neuroimaging meta-analyses

In order to examine potential biases in which papers are included in meta-analyses, we needed to have a dataset of included papers.

In order to automatically gather such a dataset, we built a heuristic method for detecting which papers are included in a meta-analysis. This method relies on the assumption that the meta-analyses will have a table listing the papers included in the meta-analysis. The method involved going through each reference and grabbing the publication year and the first author’s last name, and then using fuzzy string comparisons to match the reference to each row of the meta-analysis’ tables (i.e., Levenshtein distance thresholded at 90%). If the reference matched the contents of a row, then that reference was considered to be included in the meta-analysis and we saved the PubMed ID.

We then removed any duplicates of papers found also in PubMed Central (n = 41) and used pubget to gather the abstracts and meta-data for the ensuing list of 652 PMIDs. This was the altered version of pubget that is able to take PMIDs (the release version of pubget only takes PMCIDs).

### Collecting dataset 4) papers associated with data on NeuroVault

We started with a total of 13,396 collections on NeuroVault. Initial preprocessing included removing empty collections and those that were created by the Neuroscout account, yielding a total of 5,881 collections. Within that group, we found 896 collections with a valid DOI link to a publication in their metadata. From the remaining 4,985, we found 27 collections with a valid DOI link in the collection description. Next, we searched PubMed for articles that matched the collection’s title and found 220 additional collections linked to articles. Finally, we conducted an extensive search using pubget to find the cases where uploaded data to NeuroVault but did not link it to a publication. We performed a query and retrieved papers that mention NeuroVault in the title, abstract, keywords, and body of the articles ("*neurovault*[All Fields]"). In this way, we found additional collections linked to a valid publication for a final sample of 1,481 collections. Of these, we were able to retrieve 1,395 using pubget. This process is outlined in Figure 9.

**Figure 9.**
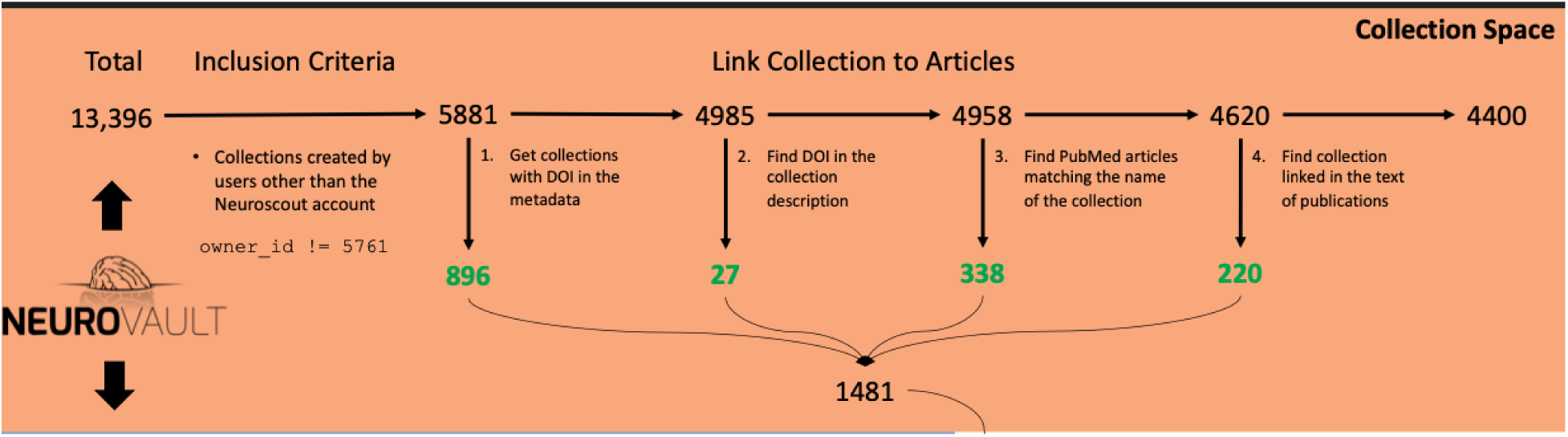
The process of obtaining articles associated with NeuroVault data.

